# The histone methyltransferase DOT1B is dispensable for stage differentiation and macrophage infection in *Leishmania mexicana*

**DOI:** 10.1101/2024.09.11.612422

**Authors:** Nicole Eisenhuth, Elisa Theres Rauh, Melina Mitnacht, Andrea Debus, Falk Butter, Ulrike Schleicher, Katerina Pruzinova, Petr Volf, Christian J. Janzen

## Abstract

Conserved histone methyltransferases of the DOT1 family are involved in replication regulation, cell cycle progression, stage differentiation and gene regulation in trypanosomatids. However, the specific functions of these enzymes depend on the host evasion strategies of the parasites. In his study, we investigated the role of DOT1B in *Leishmania mexicana*, focusing on life cycle progression and infectivity. In contrast to *Trypanosoma brucei*, in which DOT1B is essential for the differentiation of mammal-infective bloodstream forms to insect procyclic forms, *L. mexicana* DOT1B (*Lmx*DOT1B) is not critical for the differentiation of promastigotes to amastigotes *in vitro*. Additionally, there are no significant differences in the ability to infect or differentiate in macrophages or sand fly vectors between the *Lmx*DOT1B-depleted and control strains. These findings highlight the divergency of the function of DOT1B in these related parasites, suggesting genus-specific adaptations in the use of histone modifications for life cycle progression and host adaptation processes.

## INTRODUCTION

One of the greatest challenges for eukaryotic cells is to organize and maintain a dynamic genome architecture that allows precise access of protein machineries to allow DNA replication, DNA repair and gene expression. This balance between compaction and regulated access is achieved by the organization of DNA into a dynamic nucleoprotein complex called chromatin. Chromatin properties can be altered through various mechanisms, including the action of ATP-dependent chromatin remodelers, the replacement of core histones with specialized histone variants, DNA modifications, or a variety of histone modifications. Posttranslational histone modifications (PTMs) can modulate histone-histone and histone-DNA interactions, thereby altering chromatin structure. Furthermore, histone modifications can serve as binding platforms for effector proteins that mediate diverse biological processes. One of the most conserved histone modifiers is the histone methyltransferase DOT1 (<u>d</u>isruptor <u>o</u>f telomere silencing), which is the sole enzyme required for catalyzing mono-, di- and trimethylation (me1, me2 and me3) of histone H3 lysine 79 (H3K79) (Q. Feng et al., 2002; Frederiks et al., 2008; Min et al., 2003). Loss of DOT1 results in the complete absence of this methylation mark in humans, mice, flies and yeast (Deshpande et al., 2014; Y. Feng et al., 2010; Jones et al., 2008; Ng et al., 2002; Shanower et al., 2005). Initially, discovered for its role in disrupting transcriptional silencing of reporter genes adjacent to telomeres, DOT1 has multiple functions beyond telomeric silencing, including genome-wide marking of actively transcribed chromatin, cell cycle regulation, maintenance of genome integrity in response to DNA damage and initiation of replication (Farooq et al., 2016; Wood et al., 2018).

In previous work, we investigated the functions of DOT1 enzymes in trypanosomatids, which are extremely successful single-celled pathogens belonging to the class Kinetoplastida. The most significant representatives for human and domestic animal health are *Trypanosoma brucei* (African trypanosomiasis), *Trypanosoma cruzi* (Chagas disease) and *Leishmania* spp. (Leishmaniasis). Trypanosomatids have developed sophisticated cell differentiation strategies to transform into highly specialized stages to survive and proliferate in different host and vector environments. These adaptation processes involve changes in many cellular functions and are also accompanied by changes in nuclear architecture and chromatin structure (Burri et al., 1994; Povelones et al., 2012; Rout & Field, 2001; Schlimme et al., 1993). However, the exact function of chromatin structure alterations during developmental differentiation are not well understood. Already decades ago, different migration of histones in triton acid urea gels, which separate proteins according to their hydrophobicity, suggested differences in life cycle-specific histone PTMs (Burri et al., 1994; Porto et al., 2002). This observation was supported by mass spectrometry analysis comparing histone PTMs of different life cycle stages in *T. brucei* and *T. cruzi* (Janzen et al. 2006 b; Jesus et al., 2016; Mandava et al., 2007). Enzymes of the DOT1 family likely contribute to the chromatin remodeling process during differentiation. For example, *Tb*DOT1B expression is upregulated during differentiation and is essential for the differentiation of mammalian-infective bloodstream forms (BSF) to insect procyclic forms (PCF) (Dejung et al., 2016).

In contrast to yeast and humans, trypanosomes possess two paralogs of DOT1, named DOT1A and DOT1B, with distinct catalytic activities. DOT1A mediates mono- and dimethylation of H3K76, while only DOT1B also catalyzes trimethylation of the same residue (Janzen, Hake, et al., 2006). H3K76 methylation exhibits a cell cycle-dependent pattern (Gassen et al., 2012). Histones are predominantly trimethylated on H3K76 in G1-phase of the cell cycle. Newly synthesized histones are not modified during S-phase. DOT1A-mediated mono-and dimethylation can be detected at the earliest in G2 phase and seem to be crucial for replication regulation because depletion of DOT1A abolishes replication, and overexpression causes continuous reinitiation of replication (Gassen et al., 2012). In contrast, *Tb*DOT1B is not essential for cell viability under standard cell culture conditions but is involved in the repression of silent variant surface glycoprotein (VSG) genes and in switching kinetics of VSGs (Figueiredo et al., 2008; Janzen, Hake, et al., 2006). In addition to its contribution to the transcriptional control of VSG genes, *Tb*DOT1B is essential for developmental differentiation from BSF to PCF (Janzen, Hake, et al., 2006). During this process, the replicative so-called long slender forms initially differentiate into short stumpy forms, a process that is accompanied by cell cycle arrest in the G1 phase. Short stumpy forms are preadapted for survival in the midgut of the tsetse fly, and upon further environmental stimuli in the fly they are able to differentiate into the next life cycle stage, the PCF. *Tb*DOT1B-deleted cells still express the stumpy marker PAD1 and can re-enter the cell cycle during differentiation to the PCF but exhibit asymmetric nuclear division and accumulation of DNA damage after S phase (Dejung et al., 2016). Chromatin structure and nuclear architecture of BSF and PCF is different (Burri et al., 1994; Povelones et al., 2012; Rout & Field, 2001; Schlimme et al., 1993) Chromatin remodeling might occur during the first S phase after the initiation of differentiation and *Tb*DOT1B appears to be essential for this process (Dejung et al., 2016).

Little is known about the functions of DOT1 enzymes in other trypanosomatid species. In *T. cruzi*, H3K76 cell cycle-specific mono- and dimethylation patterns are conserved (Jesus et al., 2016). However, unlike in *T. brucei, Tc*DOT1B is involved in proper cell cycle progression (Nunes et al., 2020). Depletion of TcDOT1B causes a growth phenotype due to an accumulation of parasites in G2 phase and increased DNA damage. The function of TcDOT1A has not been investigated, yet. To date, little is known about DOT1 enzymes in *Leishmania*. In this study, we aimed to investigate the functions of DOT1 in *Leishmania mexicana*. We generated double-knockout cell lines of *Lmx*DOT1B and analyzed the cell cycle-dependent H3K73 methylation patterns (based on sequence alignment the homologous residue of *T. brucei* H3K76) as well as the influence of *Lmx*DOT1B on parasite life cycle progression.

Surprisingly, unlike in *T. brucei, Lmx*DOT1B is not essential for the differentiation of axenic procyclic promastigotes to amastigotes *in vitro*. Additionally, we show that *Lmx*DOT1B is dispensable to efficiently establish an infection in bone marrow-derived macrophages and sand fly vectors. Our findings suggest that while the enzymatic activity of the *Lmx*DOT1B enzyme is conserved, *L. mexicana* parasites have developed novel functions specific to their life cycle requirements.

## MATERIALS AND METHODS

### Leishmania cultivation and cell line generation

*Leishmania mexicana* procyclic promastigotes were grown in Schneider’s Drosophila medium (Serva). The medium was changed to Schneider’s Insect Medium (Sigma Aldrich) for subsequent experiments. The basic medium was supplemented with 10% heat-inactivated fetal calf serum, 10 mM 4-(2-hydroxyethyl)-1-piperazineethanesulfonic acid (HEPES), 2% sterile-filtered human urine, and 4% acid-antibiotics-pyruvate solution (AAP-mix) (200 ml AAP: 50 ml penicillin/streptomycin solution (Invitrogen), 50 ml sodium pyruvate (100 mM stock), 50 ml L-glutamine (200 mM stock), 50 ml ‘Dulbecco’s modified Eagle’s medium’ (DMEM) without phenol red with 1 g/L glucose and NaHCO3 (Sigma), 0.18 g L-asparagine, 0.58 g L-arginine)). For routine culture, *L. mexicana* procyclic promastigotes were grown at 28°C and 5% CO_2_ in humidified air. To maintain the logarithmic growth phase (1×10^5^ to 1×10^7^ cells/ml), the cells were regularly diluted based on cell counts obtained with a Coulter Counter Z2 particle counter. Transfections and drug selections were carried out as described previously (Beneke et al., 2017; Burkard et al., 2007). The transgenic Cas9 T7-expressing cell line was used as a parental cell line. To generate ΔDOT1B cells, both alleles of *Lmx*DOT1B (LmxM.20.0030) were successively replaced with a geneticin and puromycin resistance cassette following instructions on the LeishGedit website(Beneke et al., 2017). To generate the add-back cell line ΔDOT1B [DOT1B], the open reading frame of DOT1B and the flanking 3’ and 5’ UTRs were PCR-amplified and inserted into pRM005 using the EcoRI and SpeI cloning sites.

### BMDM generation and infection

All experiments followed the EU Directive 2016/63/EU, Article 23, Function A, and the German Tierschutzversuchstieranordnung (TierSchVersV, Anlage 1, Abschnitt 3). Bone marrow-derived macrophages (BMDMs) were generated from myeloid progenitor cells isolated from C57BL/6 mice. Differentiation was induced by incubation with macrophage stimulation factor (M-CSF) secreted by L929 cells in 10% conditioned medium for 7 days at 37°C and 5% CO_2_. The macrophages were harvested by incubation on ice for 5 min, gently scraped, seeded in 24-well plates with coverslips in each well and allowed to adhere to the coverslips overnight. The next day, the macrophages were infected with stationary phase promastigotes at a multiplicity of infection of 5. *Leishmania* were incubated with BMDMs for 4 h and washed before they were incubated at 37°C and 5% CO_2_ for 12 h, 24 h, 48 h or 72 h. After the designated infection time, the wells were washed with prewarmed PBS to remove dead cells. The remaining cells were fixed with 4% PFA/PBS for 10 min in the dark. After an additional three washes with PBS, the samples were permeabilized with 0.5% Triton X-100/PBS for 20 minutes. After two washing steps in 0.1% Triton X-100/PBS for 1 h, the samples were mounted on microscopy slides with DAPI Fluoromount-G mounting medium.

### Western blot analysis

Western blot analyses were carried out according to standard protocols. In brief, lysates of 2 × 10^6^ cells were separated by SDS-PAGE on 12% to 15% polyacrylamide gels and transferred to polyvinylidene difluoride (PVDF) membranes. After blocking the membranes (1 h, RT), they were incubated with primary antibodies diluted in 0.1% Tween/PBS (1 h, RT). After three washing steps with 10 ml of 0.2% Tween/PBS, the membranes were incubated with IRDye 800CW- and 680LT-coupled secondary antibodies or IRDye 680RD streptavidin (LI-COR Bioscience) in 0.1% Tween/PBS supplemented with 0.02% SDS (1 h, RT). The secondary antibodies were diluted according to the manufacturer’s instructions. The signals were imaged using a LI-COR Odyssey CLx and quantified with ImageStudio software.

### Immunofluorescence analysis

1×10^7^ cells were fixed in 1 ml of culture medium containing 4% formaldehyde for 5 min at RT. After three washes with 1 ml of PBS (1,000 g, 5 min, RT), the cells were resuspended in 300 µl of PBS. Cells were allowed to settle on poly-L-lysine-coated slides (Sigma) in a humidified chamber for 30 min. The cells were permeabilized in 100 µl of 0.2% Igepal CA-630/PBS (5 min, RT), and the slides were washed twice for 5 min in a glass slide jar filled with PBS. The cells were blocked with 100 µl of 1% BSA/PBS (1 h, RT) in a humidified chamber. After removing the blocking solution, 100 µl of primary antibody solution (anti-H3K73me2 (1:2,000) or anti-H3K73me3 (1:2,000)) in 0.1% BSA/PBS was added, and the slides were incubated for 1 h in a humidified chamber. Slides were washed three times for 5 min with PBS prior to the addition of 100 µl of secondary antibody solution (polyclonal goat Alexa Fluor 594 anti-rabbit (1:2,000) (Thermo) in 0.1% BSA/PBS supplemented with 5 μg/ml Hoechst or 1 µg/ml 4’,6-diamidino-2-phenylindole (DAPI)) or ExtrAvidinCy3 (Sigma) solution (1:100 in 0.1% BSA/PBS supplemented with 5 μg/ml Hoechst). The slides were incubated in a humidified chamber for 30 min at RT in the dark. After three washes in PBS, the cells were mounted with 10 µl of Vectashield (Vector Laboratories) and capped with coverslips. Images were captured with a Leica DMI 6000B microscope. and processed with Fiji software.

### Scanning electron microscopy

Briefly, 1×10^7^ *L. mexicana* cells per sample were harvested (1,000 g, 3 min, RT), and the supernatants were removed, except for a few microliters. The cells were fixed by the addition of 900 µl of prewarmed (27°C) Karnovsky solution (2% paraformaldehyde, 100 mM cacodylate buffer pH 7.2, 2.5% glutaraldehyde), mixed by inversion and incubated for 1 h at RT. Fixed cells were harvested (1,000 × g, 2 min, RT), washed three times with cacodylate buffer (100 mM, pH 7.2) (1,500 × g, 5 min, RT) and resuspended in 500 µl of cacodylate buffer. The attachment of cells to poly-L-lysine-coated coverslips was carried out in 24-well plates by centrifugation (1,000 g, 5 min, RT). Then, the samples were washed with 1 ml of cacodylate buffer for 5 min (1,000 g, 5 min, RT). To increase the contrast, the samples were incubated in 2% tannic acid in cacodylate buffer for 1 h at 4°C. Afterwards, the cells were washed again once with 1 ml of cacodylate buffer and three times with H_2_O for 5 min each (1,000 g, 5 min, RT). The coverslips were divided and transferred into vessels suitable for critical point drying. The samples were dehydrated in a series of ethanol (EtOH) solutions (30%, 50%, 70% and 90% EtOH for 5 min each and six times in 100% EtOH for 5 min), critical-point-dried in CO_2_, coated with gold palladium and imaged with a JEOL JSM-7500F scanning electron microscope.

### Sand fly infection

Colony of *Lutzomyia longipalpis* (Jacobina) was maintained under standard conditions as described previously (Volf & Volfova, 2011). Sand fly females were infected by feeding through a chicken skin membrane on heat-inactivated sheep blood containing 1×10^6^ log-phase promastigote *Leishmania* per ml. Engorged females were separated and maintained at 25°C with free access to 50% sugar solution. On days 2, 6 and 9 postinfection, females were dissected in drops of saline solution. The individual guts were checked for the presence of *Leishmania* promastigotes under a light microscope. The intensities of *Leishmania* infections were classified into three categories as follows (Myskova et al., 2008): light (<100 parasites/gut), moderate (100-1000 parasites/gut) and heavy (>1000 parasites/gut). The experiment was repeated twice.

## RESULTS

### The enzymatic activity of *Lmx*DOT1B is conserved

The catalytic cores of *Hs*DOT1L and *Sc*DOT1p share only a few conserved sequence motifs responsible for substrate and cofactor binding (Q. Feng et al., 2002; Min et al., 2003). Trypanosomatids possess two paralogs of DOT1 enzymes, named DOT1A and DOT1B (Janzen, Hake, et al., 2006). Both paralogs contain conserved sequence motifs responsible for substrate and cofactor binding, along with trypanosomatid-specific motifs, which confer unique product specificity and distinct functions. *Leishmania* also possesses two DOT1 paralogs (Janzen, Hake, et al., 2006) and this study focused on *Lmx*DOT1B. To determine whether *Lmx*DOT1B contains conserved motifs essential for enzymatic activity within the DOT1 family, we conducted a sequence alignment with other trypanosomal DOT1B enzymes **(Figure 1A)**. The alignment revealed conserved motifs important for co-factor and substrate binding in *Lmx*DOT1B. Specifically, *Lmx*DOT1B contains conserved motifs (I, I’, II and D2) that are crucial for cofactor S-adenosylmethionine (SAM) binding and the formation of lysin-binding channels, similar to other DOT1 enzymes (Min et al., 2003). Interestingly, in *T. brucei*, the conventional D1 motif is substituted by the CAKS sequence, which constitutes one side of the lysin-binding channel (Dindar et al., 2014). This CAKS sequence is also present in *Lmx*DOT1B. Additionally, *Lmx*DOT1B harbors the DOT1B-specific CYΦS motif (where ϕ represents hydrophobic amino acids), positioned adjacent to the CAKS sequence, contributing to the geometry of the lysine-binding channel. Furthermore, a negatively charged acidic patch near the active site of *Tb*DOT1A and *Tb*DOT1B is proposed as a binding counterpart for the positively charged residues of H3K76 and is therefore involved in nucleosome targeting (Dindar et al., 2014). This patch is conserved among trypanosomal DOT1 enzymes and is also present in *Lmx*DOT1B.

**Figure 1:**
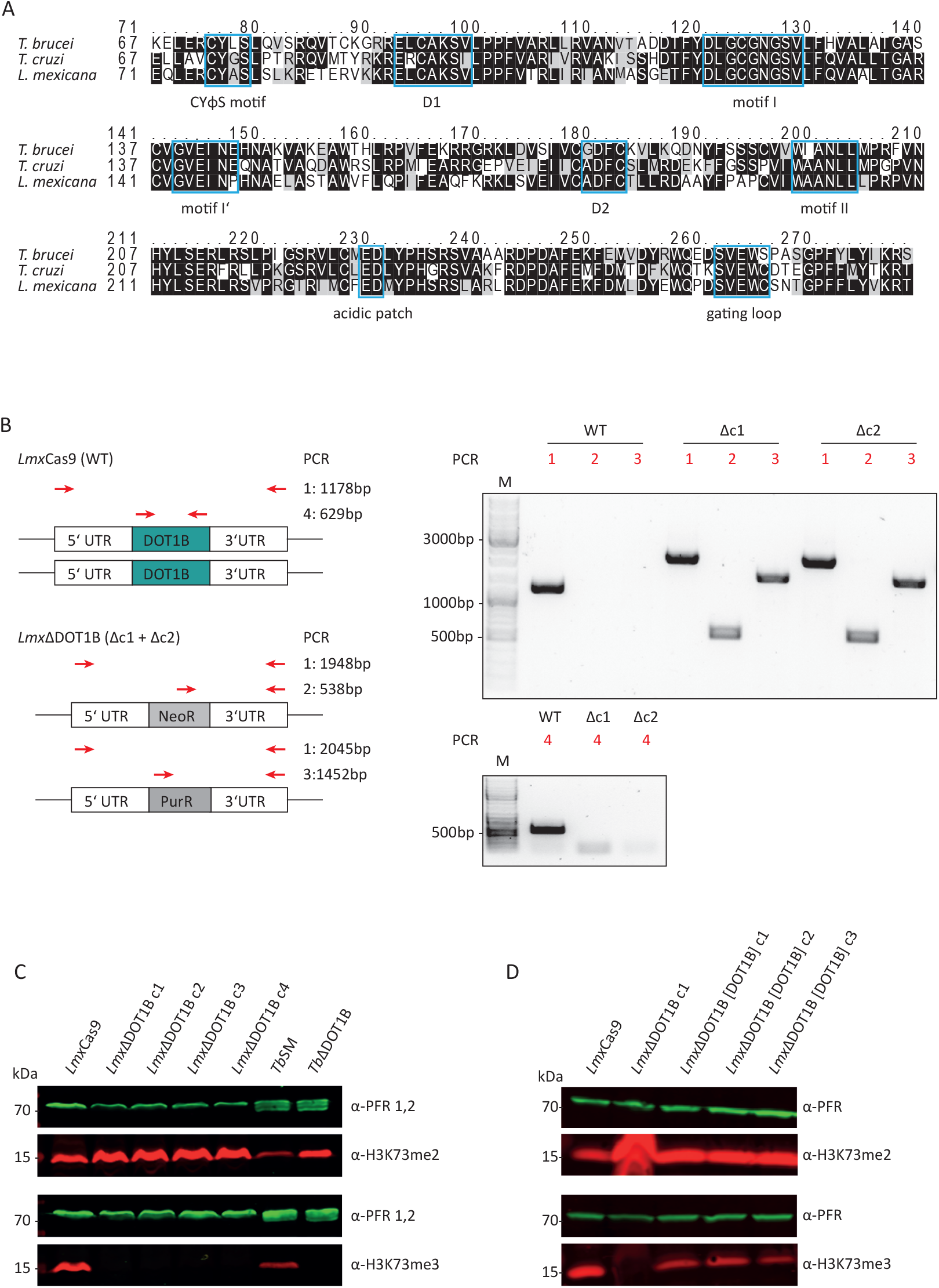
*L. mexicana* DOT1B deletion in procyclic promastigotes. **(A**) Alignments of TbDOT1B, TcDOT1B and LmxDOT1B sequences. Conserved structural motifs are highlighted with blue boxes. The black background indicates sequence identity, and the gray background indicates sequence similarity. **(B)** Schematic representation of the DOT1B gene locus in *L. mexicana* Cas9 and ΔDOT1B cells. Both alleles of DOT1B were replaced by introducing neomycin (NeoR) and puromycin (PurR) resistance cassettes (left panels). Red arrows indicate the primers used for integration control. Right panels show integration PCR with different primer pairs to verify *Lmx*DOT1B gene deletion. **(C)** Confirmation of trimethyltransferase activity in *Lmx*DOT1B. Whole-cell lysates from *Lmx*Cas9 and *Lmx*ΔDOT1B cells were subjected to immunoblotting with anti-H3K76me2 and anti-H3K76me3 antibodies as indicated. The blots were also probed with anti-PFR antibodies as a loading control. Lysates of *T. brucei* were included as controls. **(D)** Confirmation of trimethyltransferase activity in *Lmx*DOT1B through ‘add-back’ (ab) cell lines, which restored H3K73me3.

To test for conserved or novel functions of DOT1B in *L. mexicana*, DOT1B-deficient procyclic promastigote cells were generated using the CRISPR Cas9 genome editing toolkit. The *Lmx*Cas9 T7 (hereafter referred to as *Lmx*Cas9) cell line was co-transfected with two PCR-amplified sgRNA templates, creating a double-strand break upstream and downstream of the *Lmx*DOT1B ORF, thereby generating *Lmx*DOT1B knockout cells (referred to as ‘*Lmx*ΔDOT1B’). *Lmx*DOT1B deletion and successful integration of the drug resistance markers were confirmed by PCR using specific primer pairs annealing in the UTRs of *Lmx*DOT1B, drug resistance ORFs, and *Lmx*DOT1B ORFs **(Figure 1B)**. To examine the specific trimethylation activity of *Lmx*DOT1B on H3K73, whole cell lysates from *Lmx*Cas9 and *Lmx*ΔDOT1B were analyzed by Western blotting using antibodies against H3K73me3 and H3K73me2 **(Figure 1C)**. Lysates of wild-type and DOT1B-depleted *T. brucei* (*Tb*SM and *Tb*ΔDOT1B) were included as positive and negative controls, respectively. Deletion of DOT1B led to an increased dimethylation signal as shown previously, while trimethylation signals were undetectable, indicating that histone H3 is methylated at the conserved H3K73 residue in *Leishmania* and that *Lmx*DOT1B specifically catalyzes the trimethylation of H3K73. Restoring *Lmx*DOT1B expression by introducing an add-back construct (*Lmx*ΔDOT1B[DOT1B]c1-c4) also restored H3K73 trimethylation signals **(Figure 1D)** confirming that *Lmx*DOT1B is solely responsible for H3K73 trimethylation in *Leishmania mexicana*.

### The H3K73 methylation pattern is cell cycle-regulated

Precise regulation of cell cycle-dependent methylation patterns is crucial for proper replication control. *Tb*DOT1A-specific mono- and dimethylation occur only during the G2 and M phases, while trimethylation can be observed throughout the cell cycle (Gassen et al., 2012; Janzen, Hake, et al., 2006). Similar cell cycle-specific methylation patterns have been observed in *T. cruzi* (Jesus et al., 2016; Nunes et al., 2020). To investigate whether H3K73 methylation is also cell cycle-regulated in *L. mexicana*, the presence of H3K73me2 and H3K73me3 was monitored in procyclic promastigote cells using immunofluorescence analysis **(Figure 2)**. As exponentially growing promastigotes progress asynchronously through the cell cycle, the DNA of the nucleus and kinetoplast is stained to distinguish between parasites in different cell cycle phases (Wheeler et al., 2011). In *Lmx*Cas9, H3K73me2 was detected only in mitotic/cytokinetic cells, similar to the pattern observed in *T. brucei* **(Figure 2A)**. H3K73me3 was detectable throughout the cell cycle **(Figure 2B)**. In *Lmx*ΔDOT1B, no trimethylation signal was detected, confirming the loss of *Lmx*DOT1B activity. Consistent with the loss of H3K73me3 in *Lmx*ΔDOT1B, the H3K73me2 signal increased and was detectable in all cell cycle phases. These findings indicate that the DOT1-mediated cell cycle-dependent H3K76 methylation pattern is conserved in *Leishmania*.

**Figure 2:**
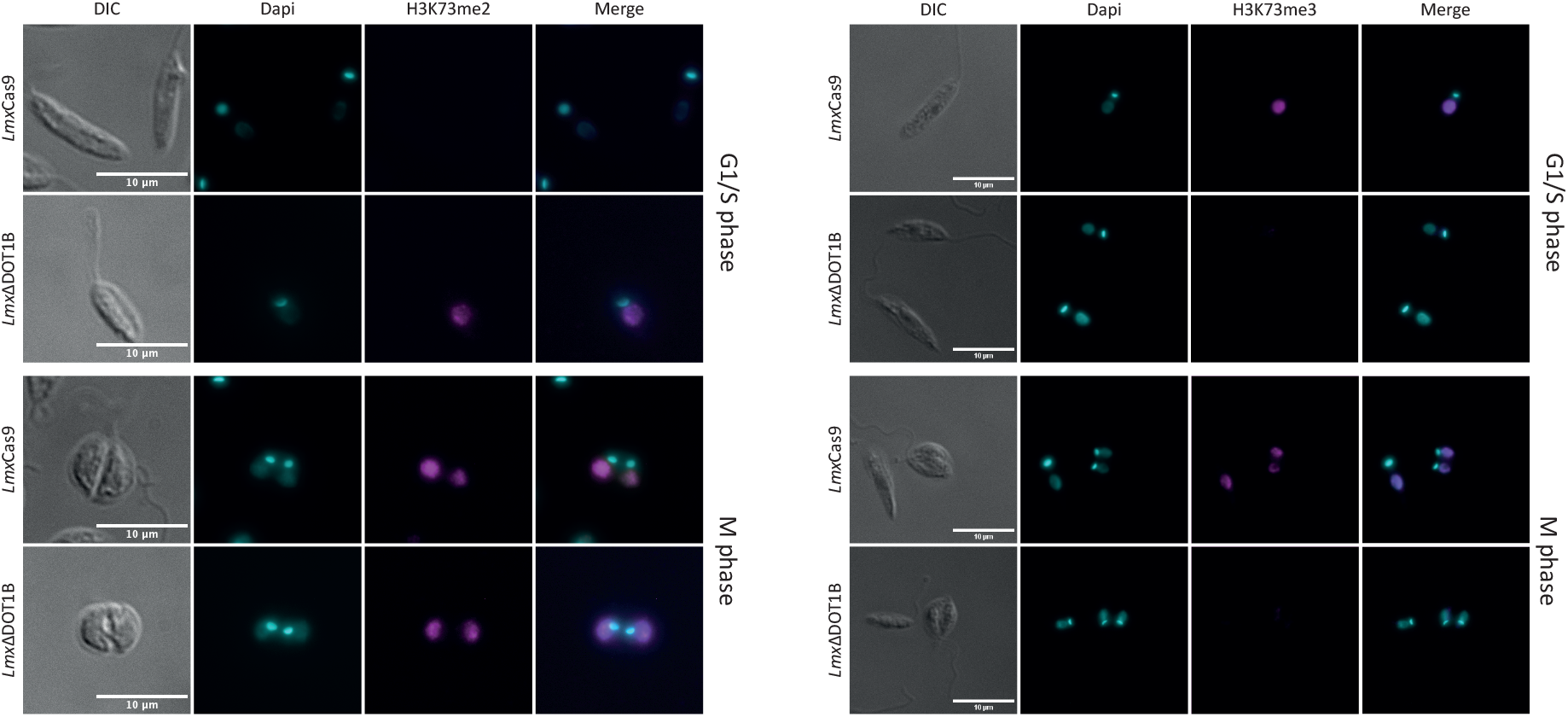
Methylation patterns in procyclic promastigotes during the cell cycle. Immunofluorescence analysis was performed to investigate the methylation patterns of H3K73 in the procyclic promastigote cells of the *Lmx*Cas9 and *Lmx*ΔDOT1B strains. H3K73me2 is predominantly observed during mitosis and cytokinesis in *Lmx*Cas9 cells. In *Lmx*ΔDOT1B cells, cell cycle specificity is absent, and H3K73me2 is also observable in G1 and S phase cells. No cell cycle-dependent distribution of H3K73me3 was detected. Notably, *Lmx*ΔDOT1B cells lack H3K73me3. DNA was stained with DAPI.

### *Lmx*DOT1B is not essential for life cycle progression

The development of a complex life cycle is fundamental for host adaptation processes of kinetoplastids. Previous studies have shown that *Tb*DOT1B is essential for differentiation from the mammalian BSF to the insect PCF (Dejung et al., 2016; Janzen, Hake, et al., 2006). In our study, the differentiation capacity of *Lmx*ΔDOT1B was compared to that of *Lmx*Cas9 **(Figure 3)**. To differentiate procyclic promastigote *Leishmania*, cells were first grown until they reached the stationary phase. For differentiation into amastigotes, stationary phase promastigotes were transferred to SDM (pH 5.4) and cultivated at elevated temperatures **(Figure 3A)**. First, we monitored growth of procyclic promastigotes in axenic culture conditions and observed a minor growth delay in DOT1B-depleted parasites (left panel). No significant differences were detected in the proportions of dead cells in either the promastigote or amastigote stage, as determined by live/dead analysis via flow cytometry **(Supplementary Figure 1)**. Thus, cell death was ruled out as a cause of the slight growth delay in the *Lmx*ΔDOT1B cells. DOT1B-depleted parasites reached the stationary phase with a slight delay but at identical cell density compared to WT cells (middle panel). Finally, *Lmx*ΔDOT1B and WT cells were differentiated to amastigotes *in vitro*. Both cell lines started to proliferate after induction of differentiation, although *Lmx*ΔDOT1B parasites again exhibited a slight growth delay (left panel). The transition from the procyclic to the amastigote stage in *L. mexicana* is marked by profound morphological changes. While procyclic cells exhibit an elongated shape and long flagellum, amastigote cells are ovoid-shaped and possess short flagella. We used scanning electron microscopy to compare morphological changes in *Lmx*ΔDOT1B and WT cells during the differentiation process **(Figure 3B)**. No discernible differences were observed between the two populations, which led us to conclude that DOT1B is not involved in the morphological transition during differentiation of *L. mexicana*. In addition to morphology, the differentiation process can be further monitored by the analysis of stage-specific protein abundancy. For example, the paraflagellar rod protein (PFR) is downregulated during differentiation to amastigotes due to the shortening of the flagellum. Thus, we measured PFR expression levels via Western blot analysis and evaluated different life cycle stages in parental *Lmx*Cas9 cells and *Lmx*ΔDOT1B cells **(Figure 3C)**. No major differences could be detected between WT and DOT1B-depleted parasites. While the PFR signal is present in lysates of procyclic and stationary phase promastigotes, it is undetectable in amastigotes for both *Lmx*Cas9 and *Lmx*ΔDOT1B cells.

**Figure 3:**
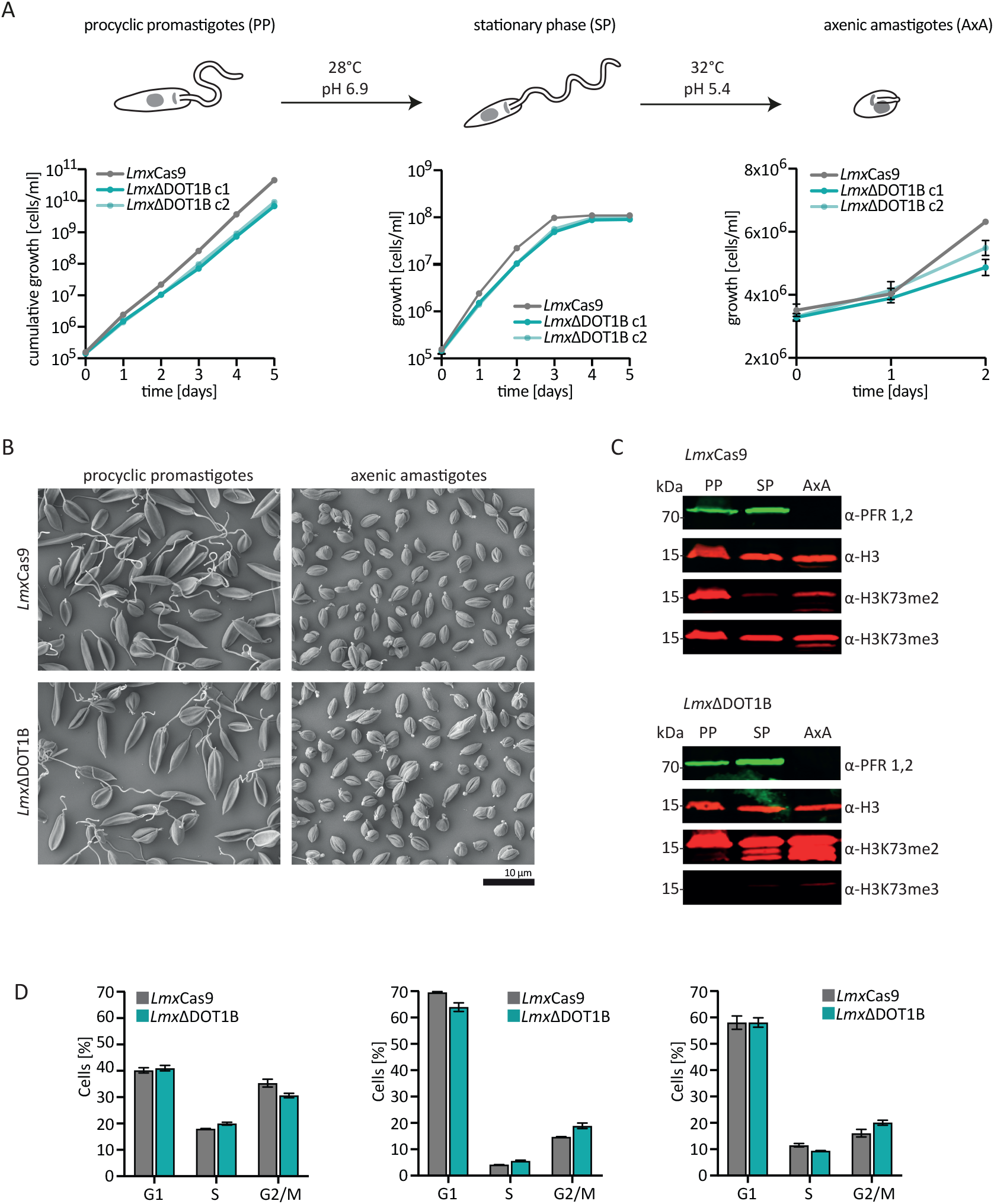
*In vitro* differentiation of *Lmx*ΔDOT1B. **(A)**. For differentiation, logarithmically growing procyclic promastigotes were diluted to 1×105 cells/ml and left to grow to stationary phase for five days (28°C, pH 6.9). Stationary phase promastigotes were differentiated into axenic amastigotes at 32°C and pH 5.4. The growth of two *Lmx*ΔDOT1B clones and *Lmx*WT cells was monitored during the differentiation process. Compared to the *Lmx*WT cells, the *Lmx*ΔDOT1B mutants exhibited slightly slower growth. The means and standard deviations of four biological replicates are shown. **(B)** Procyclic promastigote and axenic amastigote *Lmx*ΔDOT1B populations show life cycle stage-specific morphologies, as observed by SEM. *Lmx*WT cells were analyzed as a control. **(C)** Western blots confirming the expression of the promastigote-specific marker PFR in procyclic promastigotes (PPs) and its loss in axenic amastigotes (AxAs). The H3K73me2 signal is undetectable in G1-arrested stationary phase (SP) WT cells but reoccurs in AxA cells upon resumption of the cell cycle. *Lmx*ΔDOT1B results in a decrease in H3K73 trimethylation and a simultaneous increase in dimethylation, irrespective of the life cycle stage. Histone H3 was used as a protein loading control. **(D)** Evaluation of the cell cycle profiles of fixed PI-stained *Lmx*Cas9 and *Lmx*ΔDOT1B cells at different life cycle stages. Growth to the stationary phase leads to the arrest of cells in the G1 phase. Following differentiation into the amastigote form, the cells resume their cell cycle.

The absence of a H3K73me2 in stationary phase *Lmx*Cas9 points toward cell cycle arrest in the G1 phase, since we previously demonstrated that the H3K73me2 mark is missing in this phase. To determine whether DOT1B-depletion influences cell cycle progression during differentiation in *L. mexicana*, cells were stained with propidium iodide (PI) and analyzed by flow cytometry **(Figure 3D; Supplementary Figure 2)**. The fluorescence intensity of the DNA-intercalating PI correlated with the DNA content of the cells, allowing us to classify the cells into G1-, S- and G2/M phase populations. When comparing parental with DOT1B KO cells, no statistically significant differences were observed. Forty percent of the exponentially growing procyclic promastigote WT cells could be assigned to G1 phase, 20% to S phase and 35% to G2 phase. The proportion of G1-phase cells increased to 70% with a concomitant decrease in the other cell cycle in stationary procyclic promastigotes, demonstrating that most cells were arrested in the G1 phase. After differentiation to the amastigote form, both cell lines slowly resumed their cell cycle, which was reflected in decreasing G1- and increasing S- and G2-phase populations.

### Loss of LmxDOT1B does not impair sand fly and macrophage infection

Next, we addressed the ability of WT and DOT1B KO *Leishmania* to infect the sand fly vector *L. longipalpis*. The female sand flies were infected by feeding heat-inactivated sheep blood containing log phase promastigote *Lmx*Cas9 or *Lmx*ΔDOT1B *Leishmania*. Engorged females were separated, dissected at 2, 6 or 9 days post infection and evaluated microscopically according to their infection rates and intensities. To monitor the progress of infection, the localization of the parasites within the sand fly midgut was also assessed.

No significant differences in infection rates or intensities were detected between *Lmx*Cas9 and *Lmx*ΔDOT1B. Both experimental groups showed similar proportions of light, moderate and heavy infection intensities **(Figure 4A)** and yielded infection rates of approximately 80% throughout the experiment. The course of infection was also quite similar between *Lmx*Cas9 and *Lmx*ΔDOT1B. Starting on day 6, in 70-80% of infected sand fly females, *Leishmania* parasites colonize the stomodeal valve, an important prerequisite for further transmission during the parasite’s life cycle **(Figure 4B)**. These data suggest that *Lmx*DOT1B is dispensable for an efficient and transmissible infection of the sand fly vector.

**Figure 4:**
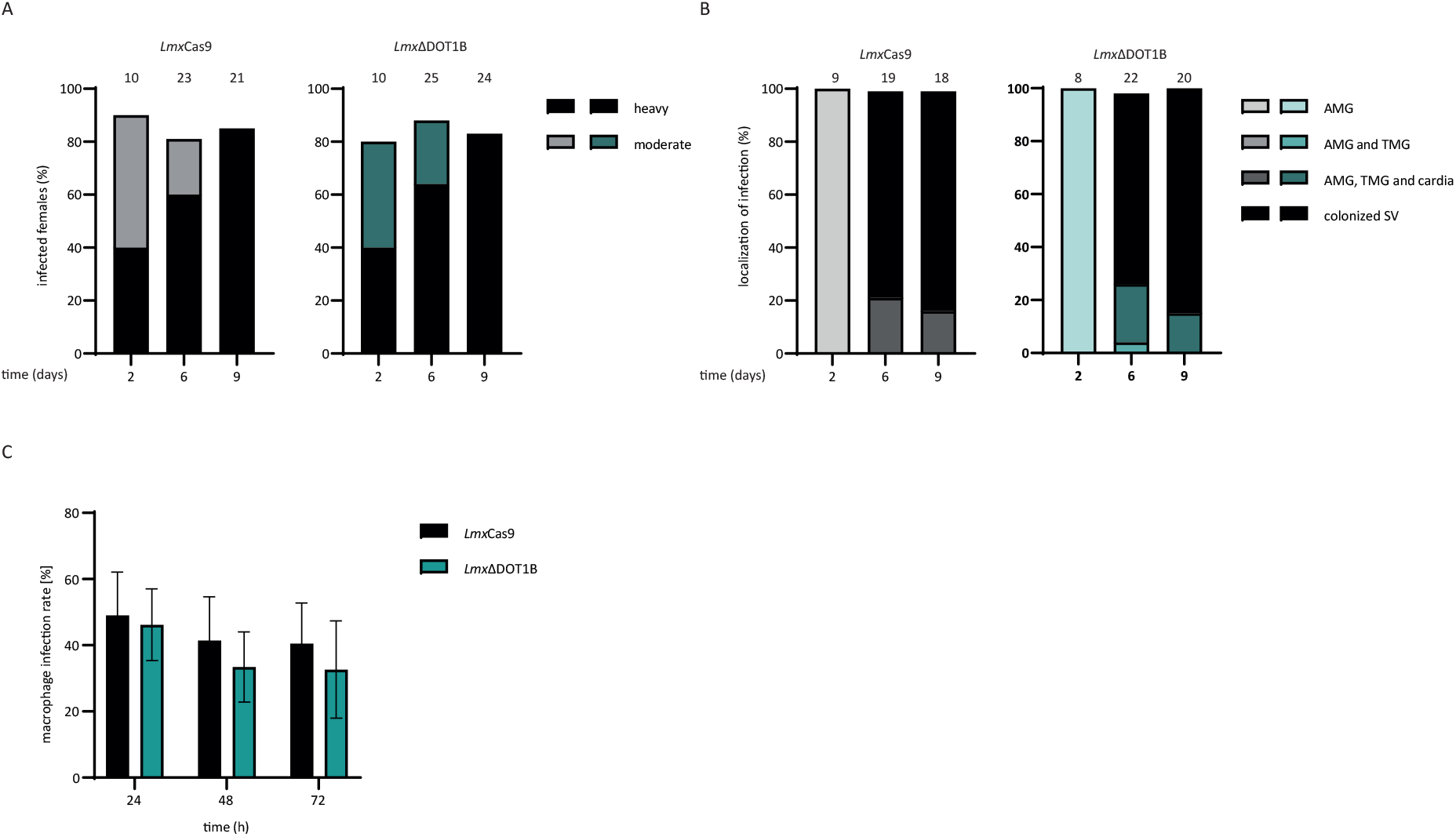
Infection of sand flies and macrophages. **(A)** *Lmx*ΔCas9 and *Lmx*ΔDOT1B infection rates and intensities in *Lutzomyia longipalpis* were assessed on days 2, 6, and 9 post-infection of the sand flies. The numbers above the bars indicate the total number of dissected females. Leishmania infection intensities were categorized into three classes: light (< 100 parasites/gut), moderate (100-1000 parasites/gut) and heavy (>1000 parasites/gut). **(B)** Localization of infections in the abdominal midgut (AMG), thoracic midgut (TMG), and stomodeal valve (SV) on days 2, 6, and 9 postinfection. The numbers above the bars represent the total number of evaluated females with detectable infections. **(C)** Stationary phase promastigotes were used to infect C57BL/6 BMDMs at a multiplicity of infection of 5. The percentage of infected macrophages was assessed at 4, 12, 24, 48 and 72 h post infection. The mean and standard deviation of three replicate experiments are shown. For each infection experiment, 200 macrophages were analyzed.

An important part of the *Leishmania* life cycle is the ability to infect mammalian hosts, where the parasites differentiate and multiply inside macrophages. They developed sophisticated strategies to modulate the host immune response to survive within a hostile environment, which is usually responsible for the elimination of pathogens. To determine whether such complex processes are impaired upon the loss of *Lmx*DOT1B, we infected bone marrow-derived macrophages (BMDMs) with *Lmx*ΔDOT1B parasites to analyze the uptake and survival rates of the mutant parasites within the first three days post infection. *Lmx*DOT1B parasites showed no statistically significant differences in the ability to infect macrophages compared to *Lmx*Cas9 cells **(Figure 4C)**. Both cell lines were able to achieve approximately 50% initial infection rates, which decline to approximately 40% 72 h post infection.

In summary, in contrast to other Kinetoplastida, *Lmx*DOT1B is not essential for normal growth in cell culture or stage differentiation. Furthermore, it is dispensable to establish an infection efficiently in mammalian macrophages and in the insect vector.

## Discussion

*T. brucei, T. cruzi*, and *L. mexicana*, despite all being members of the kinetoplastida, exhibit distinct dependencies on DOT1B for their growth, stage development and pathogenicity. Understanding these differences at a molecular level can provide valuable insights into parasite biology and open possibilities for targeted therapeutic strategies that exploit these species-specific dependencies. In this study, we aimed to shed light on the function of DOT1B in *L. mexicana*.

In *T. brucei* and *T. cruzi*, DOT1B is necessary for some of these processes. Although *Tb*DOT1B is not essential for cell viability under standard cell culture conditions, it is involved in the repression of silent variant surface glycoprotein genes and in the kinetics of VSG switching (Figueiredo et al., 2008; Janzen, Hake, et al., 2006). Furthermore, *Tb*DOT1B is crucial for developmental differentiation from bloodstream forms to procyclic forms (Dejung et al., 2016; Janzen, Hake, et al., 2006). *Tb*DOT1B knockout cells, differentiation to so-called stumpy forms, including G1 arrest is, not impaired. However, after re-entry into the cell cycle, ΔDOT1B *T. brucei* exhibit defects in karyokinesis and accumulation of DNA damage. This suggests that *Tb*DOT1B is required for changes in chromatin structure that occur in the first S-phase after re-entry into cell cycle progression during developmental differentiation.

In *T. cruzi*, DOT1B also plays a significant role in cell cycle progression. Deletion of *Tc*DOT1B causes a growth phenotype characterized by an accumulation of parasites in the G2 phase and increased DNA damage markers. While the exact mechanism by which DOT1B is necessary for normal cell cycle progression in *T. cruzi* remains elusive, Nuenes and colleagues suggested that DOT1B is involved in checkpoint activation after DNA damage accumulation. (Nunes et al., 2020)

The function of DOT1B in *Leishmania* seems to be different compared to the aforementioned trypanosomes. Our findings demonstrate that DOT1B is dispensable for the parasite’s life cycle progression, infectivity, and differentiation. However, as in *T. brucei* and *T. cruzi*, dimethylation could be observed only in mitotic or cytokinetic cells, while trimethylation was detectable throughout the cell cycle, indicating that LmxDOT1-mediated methylation could also be involved in DNA replication regulation as described in other trypanosomes (Gassen et al., 2012). The mechanisms of DNA replication and damage repair in *Leishmania spp*. are not well understood. One explanation for the absence of defects in cell cycle progression and karyokinesis after DOT1B depletion might be that *Leishmania* parasites mainly utilize different strategies for DNA damage detection compared to other kinetoplastids and rearrangements of chromatin structure might not be necessary during stage development in these parasites.

## Funding

KP and PV were supported by European Regional Development Fund (ERDF) - project CePaViP 16_019/0000759. NE and CJJ were supported by Deutsche Forschungsgemeinschaft (JA 1013/7-1)

## Conflict of interest

The authors declare that the research was conducted without commercial or financial relationships that could be construed as potential conflicts of interest.

## Notes

### Competing Interest Statement

The authors have declared no competing interest.

## References

Beneke, T., Madden, R., Makin, L., Valli, J., Sunter, J., & Gluenz, E. (2017). A CRISPR Cas9 high-throughput genome editing toolkit for kinetoplastids. Royal Society Open Science, 4(5), 170095. 10.1098/rsos.170095

Burkard, G., Fragoso, C. M., & Roditi, I. (2007). Highly efficient stable transformation of bloodstream forms of Trypanosoma brucei. Molecular and Biochemical Parasitology, 153(2), 220–223. 10.1016/j.molbiopara.2007.02.008

Burri, M., Schlimme, W., Betschart, B., & Hecker, H. (1994). Characterization of the histones of Trypanosoma brucei brucei bloodstream forms. Acta Tropica, 58(3–4), 291–305. 10.1016/0001-706x(94)90023-x

Dejung, M., Subota, I., Bucerius, F., Dindar, G., Freiwald, A., Engstler, M., Boshart, M., Butter, F., & Janzen, C. J. (2016). Quantitative Proteomics Uncovers Novel Factors Involved in Developmental Differentiation of Trypanosoma brucei. PLoS Pathogens, 12(2), e1005439. 10.1371/journal.ppat.1005439

Deshpande, A. J., Deshpande, A., Sinha, A. U., Chen, L., Chang, J., Cihan, A., Fazio, M., Chen, C., Zhu, N., Koche, R., Dzhekieva, L., Ibáñez, G., Dias, S., Banka, D., Krivtsov, A., Luo, M., Roeder, R. G., Bradner, J. E., Bernt, K. M., & Armstrong, S. A. (2014). AF10 Regulates Progressive H3K79 Methylation and HOX Gene Expression in Diverse AML Subtypes. Cancer Cell, 26(6), 896–908. 10.1016/j.ccell.2014.10.009

Dindar, G., Anger, A. M., Mehlhorn, C., Hake, S. B., & Janzen, C. J. (2014). Structure-guided mutational analysis reveals the functional requirements for product specificity of DOT1 enzymes. Nature Communications, 5(1), 5313. 10.1038/ncomms6313

Farooq, Z., Banday, S., Pandita, T. K., & Altaf, M. (2016). The many faces of histone H3K79 methylation. Mutation Research/Reviews in Mutation Research, 768, 46–52. 10.1016/j.mrrev.2016.03.005

Feng, Q., Wang, H., Ng, H. H., Erdjument-Bromage, H., Tempst, P., Struhl, K., & Zhang, Y. (2002). Methylation of H3-Lysine 79 Is Mediated by a New Family of HMTases without a SET Domain. Current Biology, 12(12), 1052–1058. 10.1016/s0960-9822(02)00901-6

Feng, Y., Yang, Y., Ortega, M. M., Copeland, J. N., Zhang, M., Jacob, J. B., Fields, T. A., Vivian, J. L., & Fields, P. E. (2010). Early mammalian erythropoiesis requires the Dot1L methyltransferase. Blood, 116(22), 4483–4491. 10.1182/blood-2010-03-276501

Figueiredo, L. M., Janzen, C. J., & Cross, G. A. M. (2008). A Histone Methyltransferase Modulates Antigenic Variation in African Trypanosomes. PLoS Biology, 6(7), e161. 10.1371/journal.pbio.0060161

Frederiks, F., Tzouros, M., Oudgenoeg, G., Welsem, T.van, Fornerod, M., Krijgsveld, J., & Leeuwen, F.van. (2008). Nonprocessive methylation by Dot1 leads to functional redundancy of histone H3K79 methylation states. Nature Structural & Molecular Biology, 15(6), 550–557. 10.1038/nsmb.1432

Gassen, A., Brechtefeld, D., Schandry, N., Arteaga-Salas, J. M., Israel, L., Imhof, A., & Janzen, C. J. (2012). DOT1A-dependent H3K76 methylation is required for replication regulation in Trypanosoma brucei. Nucleic Acids Research, 40(20), 10302–10311. 10.1093/nar/gks801

Janzen, C. J., Fernandez, J. P., Deng, H., Diaz, R., Hake, S. B., & Cross, G. A. M. (2006). Unusual histone modifications in Trypanosoma brucei. FEBS Letters, 580(9), 2306–2310. 10.1016/j.febslet.2006.03.044

Janzen, C. J., Hake, S. B., Lowell, J. E., & Cross, G. A. M. (2006). Selective Di-or Trimethylation of Histone H3 Lysine 76 by Two DOT1 Homologs Is Important for Cell Cycle Regulation in Trypanosoma brucei. Molecular Cell, 23(4), 497–507. 10.1016/j.molcel.2006.06.027

Jesus, T. C. L. de, Nunes, V. S., Lopes, M. de C., Martil, D. E., Iwai, L. K., Moretti, N. S., Machado, F. C., Lima-Stein, M.L.de, Thiemann, O. H., Elias, M. C., Janzen, C., Schenkman, S., & Cunha, J. P. C.da. (2016). Chromatin Proteomics Reveals Variable Histone Modifications during the Life Cycle of Trypanosoma cruzi. Journal of Proteome Research, 15(6), 2039–2051. 10.1021/acs.jproteome.6b00208

Jones, B., Su, H., Bhat, A., Lei, H., Bajko, J., Hevi, S., Baltus, G. A., Kadam, S., Zhai, H., Valdez, R., Gonzalo, S., Zhang, Y., Li, E., & Chen, T. (2008). The Histone H3K79 Methyltransferase Dot1L Is Essential for Mammalian Development and Heterochromatin Structure. PLoS Genetics, 4(9), e1000190. 10.1371/journal.pgen.1000190

Mandava, V., Fernandez, J. P., Deng, H., Janzen, C. J., Hake, S. B., & Cross, G. A. M. (2007). Histone modifications in Trypanosoma brucei. Molecular and Biochemical Parasitology, 156(1), 41–50. 10.1016/j.molbiopara.2007.07.005

Min, J., Feng, Q., Li, Z., Zhang, Y., & Xu, R.-M. (2003). Structure of the Catalytic Domain of Human DOT1L, a Non-SET Domain Nucleosomal Histone Methyltransferase. Cell, 112(5), 711–723. 10.1016/s0092-8674(03)00114-4

Myskova, J., Votypka, J., & Volf, P. (2008). Leishmania in Sand Flies: Comparison of Quantitative Polymerase Chain Reaction with Other Techniques to Determine the Intensity of Infection. Journal of Medical Entomology, 45(1), 133–138. 10.1093/jmedent/45.1.133

Ng, H. H., Feng, Q., Wang, H., Erdjument-Bromage, H., Tempst, P., Zhang, Y., & Struhl, K. (2002). Lysine methylation within the globular domain of histone H3 by Dot1 is important for telomeric silencing and Sir protein association. Genes & Development, 16(12), 1518–1527. 10.1101/gad.1001502

Nunes, V. S., Moretti, N. S., Silva, M.S.da, Elias, M. C., Janzen, C. J., & Schenkman, S. (2020). Trimethylation of histone H3K76 by Dot1B enhances cell cycle progression after mitosis in Trypanosoma cruzi. Biochimica et Biophysica Acta (BBA) - Molecular Cell Research, 1867(7), 118694. 10.1016/j.bbamcr.2020.118694

Porto, R. M., Amino, R., Elias, M. C. Q., Faria, M., & Schenkman, S. (2002). Histone H1 is phosphorylated in non-replicating and infective forms of Trypanosoma cruzi. Molecular and Biochemical Parasitology, 119(2), 265–271. 10.1016/s0166-6851(01)00430-3

Povelones, M. L., Gluenz, E., Dembek, M., Gull, K., & Rudenko, G. (2012). Histone H1 Plays a Role in Heterochromatin Formation and VSG Expression Site Silencing in Trypanosoma brucei. PLoS Pathogens, 8(11), e1003010. 10.1371/journal.ppat.1003010

Rout, M. P., & Field, M. C. (2001). Isolation and Characterization of Subnuclear Compartments from Trypanosoma brucei IDENTIFICATION OF A MAJOR REPETITIVE NUCLEAR LAMINA COMPONENT*. Journal of Biological Chemistry, 276(41), 38261–38271. 10.1074/jbc.m104024200

Schlimme, W., Burri, M., Bender, K., Betschart, B., & Hecker, H. (1993). Trypanosoma brucei brucei: differences in the nuclear chromatin of bloodstream forms and procyclic culture forms. Parasitology, 107(3), 237–247. 10.1017/s003118200007921x

Shanower, G. A., Muller, M., Blanton, J. L., Honti, V., Gyurkovics, H., & Schedl, P. (2005). Characterization of the grappa Gene, the Drosophila Histone H3 Lysine 79 Methyltransferase. Genetics, 169(1), 173–184. 10.1534/genetics.104.033191

Volf, P., & Volfova, V. (2011). Establishment and maintenance of sand fly colonies. Journal of Vector Ecology, 36(1), S1–S9. 10.1111/j.1948-7134.2011.00106.x

Wheeler, R. J., Gluenz, E., & Gull, K. (2011). The cell cycle of Leishmania: morphogenetic events and their implications for parasite biology. Molecular Microbiology, 79(3), 647–662. 10.1111/j.1365-2958.2010.07479.x

Wood, K., Tellier, M., & Murphy, S. (2018). DOT1L and H3K79 Methylation in Transcription and Genomic Stability. Biomolecules, 8(1), 11. 10.3390/biom8010011

