## Supplementary Figures for "The histone methyltransferase DOT1B is dispensable for stage differentiation and macrophage infection in *Leishmania mexicana*"

Supplementary Figure 1

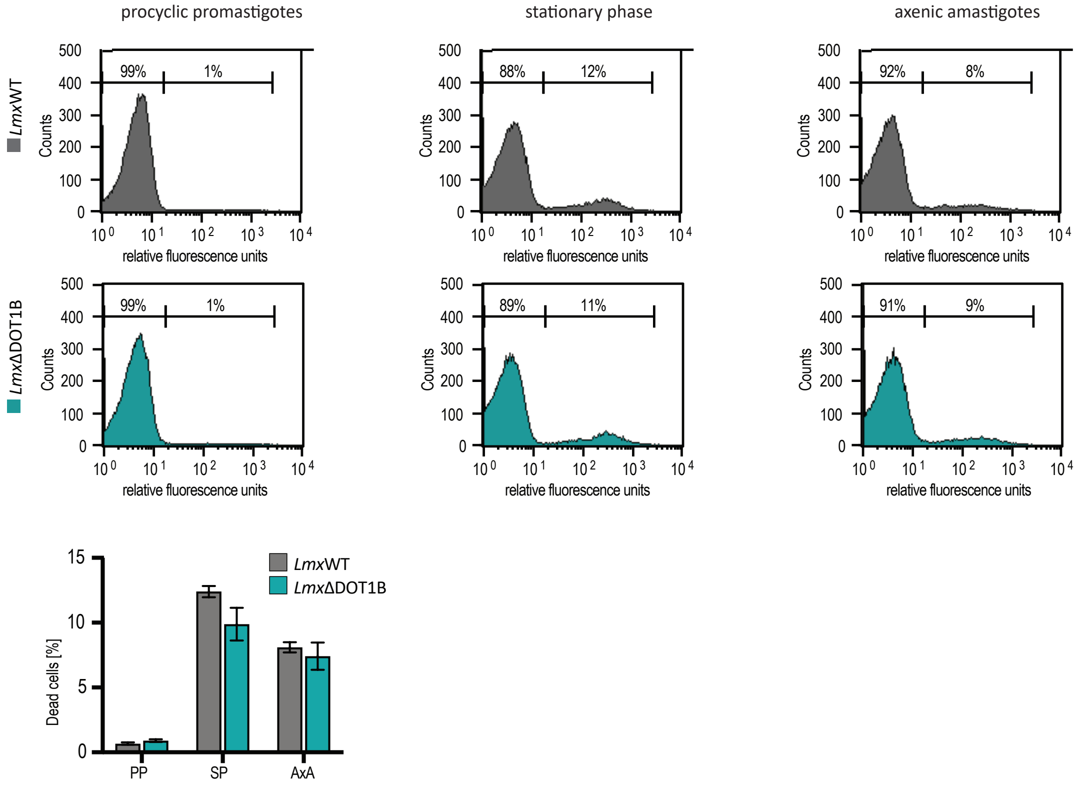


**Figure S1: Growth delay of LmxΔDOT1B cells is not caused by dying cells.**
Flow cytometry analysis of PI-stained LmxΔDOT1B cells in the different life cycle stages. Left gate shows the living cells, defined by the *Lmx*WT cells. The right gate shows dead cells, which were defined by *Lmx*WT cells after treatment with blasticidin (BLAS). One representative replicate is shown. The mean percentage (n=3) of dead cells of each life cycle stage is displayed in the bar graph (PP (procyclic promastigotes); SP (stationary phase); AxA (axenic amastigotes)).

Supplementary Figure 2

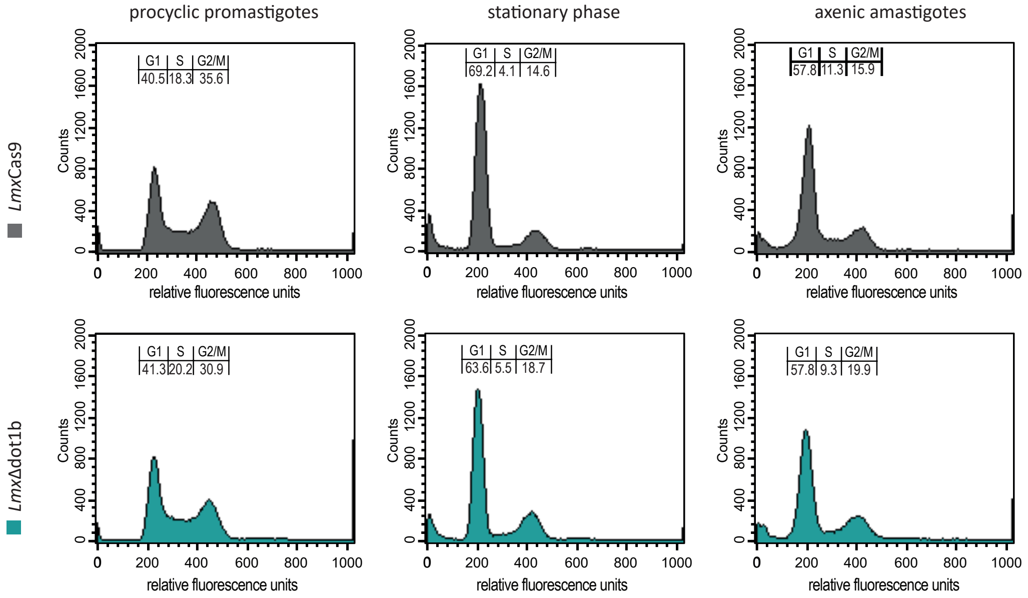


**Figure S2: Cell cycle profiles during differentiation of LmxΔDOT1B cells.**
Cell cycle profiles of fixed PI-stained *Lmx*ΔDOT1B cells in the different life cycle stages. *Lmx*WT cells served as a control. One representative replicate is shown. The percentage of the mean values of each cell cycle stage is displayed in the bar graph in figure 4 (n=3).
